## Supplementary Information for "Modeling microbial cross-feeding at intermediate scale portrays community dynamics and species coexistence"

June 4, 2020

**2 Supplementary Tables** **17**

**3 Supplementary Figures** **22**

**4 Supplementary References** **33**

### 1 Supplementary Texts

#### 1.1 A Biophysical Modeling Framework for Microbial Community

In this study, we present a biophysical model that combines population dynamics of different cell types that complete for external nutrients and their intracellular metabolism. The population dynamics part of our model (i.e., cell population expansion and contraction) is similar to previous models [1, 2, 3] but the part with respect to the intracellular metabolism is very different. We considered the internal metabolism of nutrient uptake and conversion via a simplified 3-level metabolic network that captures resource transformation from growth substrates to metabolic building blocks and then to biomass. Growth substrates, either substitutable or non-substitutable, are molecules that are able to support cell growth as the sole sources that supply nutrients they contain. Metabolic building blocks are incorporated into different functional units of biomass and thus assume to be non-substitutable. Note that in our coarse-grained picture, building blocks can be any metabolite intermediates that serve as precursors of molecules that are actually integrated into biomass. In some cases, a molecule can be a growth substrate for one cell type and a metabolite precursor for another.

The variables we considered in our model include: (1) Density of active cells of $n_c$ microbial populations with distinct cell types ($N_l$, $l = 1, 2, \cdots, n_c$); (2) Concentration of $n_s$ growth substrates in the culture medium ($S_i$, $i = 1, 2, \cdots, n_s$) and in cell type $l$ ($\hat{S}_{l,i}$, $i = 1, 2, \cdots, n_s$); (3) Concentration of $n_m$ metabolic building blocks in the culture medium ($M_j$, $j = 1, 2, \cdots, n_m$) and in cell type $l$ ($\hat{M}_{l,j}$, $j = 1, 2, \cdots, n_m$). The biochemical reactions that define the relationships among these variables are given below

$$S_i \xrightarrow{J_{l,i}^{upt,S}} \hat{S}_{l,i} \quad l = 1, 2, \cdots, n_c, \; i = 1, 2, \cdots, n_s \tag{1}$$

$$M_j \xrightarrow{J_{l,j}^{upt,M}} \hat{M}_{l,j} \quad l = 1, 2, \cdots, n_c, \; j = 1, 2, \cdots, n_m \tag{2}$$

$$\sum_{i=1}^{n_s} (1/\delta_{l,i,j}) \hat{S}_{l,i} \xrightarrow{J_{l,j}^{syn}} \hat{M}_{l,j} \quad l = 1, 2, \cdots, n_c, \; j = 1, 2, \cdots, n_m \tag{3}$$

$$\hat{M}_{l,j} \xrightarrow{J_{l,j}^{leak}} M_j \quad l = 1, 2, \cdots, n_c, \; j = 1, 2, \cdots, n_m \tag{4}$$

$$\sum_{i=1}^{n_s} (1/\gamma_{l,i}) \hat{S}_{l,i} + \sum_{j=1}^{n_m} (1/\gamma_{l,j}) \hat{M}_{l,j} \xrightarrow{J_{l}^{grow}} \text{An active cell of population } l \quad l = 1, 2, \cdots, n_c \tag{5}$$

$$\text{An active cell of population } l \xrightarrow{J_{l}^{death}} \text{An inactive cell of population } l \quad l = 1, 2, \cdots, n_c \tag{6}$$

$$\hat{S}_{l,i} \xrightarrow{J_{l,i}^{sink,S}} \emptyset \quad l = 1, 2, \cdots, n_c, \; i = 1, 2, \cdots, n_s \tag{7}$$

$$\hat{M}_{l,j} \xrightarrow{J_{l,j}^{sink,M}} \emptyset \quad l = 1, 2, \cdots, n_c, \; j = 1, 2, \cdots, n_m \tag{8}$$

where $J$'s represents reaction rates (fluxes). Equation (1) and (2) describe resource uptake into intracellular space. Equation (3) describes biosynthesis of intermediate metabolites $\hat{M}_{l,j}$ from substrates $\hat{S}_{l,i}$, where $1/\delta_{l,i,j}$ is the number of molecules of substrate $\hat{S}_i$ consumed for every one molecule of metabolite $\hat{M}_j$ produced by cell type $l$. Equation (4) describes the leakage of intracellular metabolites $\hat{M}_{l,j}$ into the environment. Equation (5) describes biomass synthesis from both substrate $\hat{S}_{l,i}$ and metabolic building blocks $\hat{M}_{l,j}$, where $\gamma_{l,i}$ and $\gamma_{l,j}$ are biomass yield coefficients (alternatively, $1/\gamma_{l,i}$ and $1/\gamma_{l,j}$ represent the numbers of molecules of $\hat{S}_{l,i}$ and $\hat{M}_{l,j}$ needed to construct one active cell of cell type $l$ respectively). Importantly, the first sum over substrates in Equation (5) approximates biomass composition from other building blocks that are not explicitly modeled. Equation (6) describes a decrease in viable biomass, i.e., cell death. Equation (7) and (8) represent sink reactions that consume substrates $\hat{S}_{l,i}$ and metabolites $\hat{M}_{l,j}$ through non-growth metabolic processes and degradation pathways.

We consider constant supply rate of growth substrates in a chemostat culture with dilution rate $D$. The eight reactions shown above can be translated to the following differential equations

$$\frac{d[S_i]}{dt} = D(S_{0,i} - [S_i]) - \sum_{l=1}^{n_c} J_{l,i}^{upt,S} N_l \qquad i = 1, 2, \cdots, n_s \tag{9}$$

$$\frac{dN_l}{dt} = N_l \left( J_l^{grow} - J_l^{death} - D \right) \qquad l = 1, 2, \cdots, n_c \tag{10}$$

$$\frac{d[M_j]}{dt} = D(M_{0,j} - [M_j]) + \sum_{l=1}^{n_c} \left( J_{l,j}^{leak} - J_{l,j}^{upt,M} \right) N_l \qquad j = 1, 2, \cdots, n_m \tag{11}$$

$$\frac{d[\hat{S}_{l,i}]}{dt} = J_{l,i}^{upt,S} - \sum_{j=1}^{n_m} (1/\delta_{l,i,j}) J_{l,j}^{syn} - (1/\gamma_{l,i}) J_l^{grow} - J_{l,i}^{sink,S} \tag{12}$$

$$\frac{d[\hat{M}_{l,j}]}{dt} = J_{l,j}^{syn} + J_{l,j}^{upt,M} - J_{l,j}^{leak} - (1/\gamma_{l,j}) J_l^{grow} - J_{l,j}^{sink,M} \tag{13}$$

where $[\cdots]$ denotes molecular concentration. $S_{0,i}$ and $M_{0,j}$ are the concentrations of $S_i$ and $M_j$ in the chemostat feed medium respectively.

Bacterial growth occurs at much faster time scale compared to that associated with intracellular metabolic processes [3]. It is therefore reasonable to separate time scales by assuming quasi-steady-state for intracellular growth substrates $\hat{S}_{l,i}$ and metabolic building blocks $\hat{M}_{l,j}$. Solving Equation (12) and (13) with $d[\hat{S}_{l,i}]/dt = d[\hat{M}_{l,j}]/dt = 0$ yields the internal flux-balance equations

$$J_{l,i}^{upt,S} - \sum_{j=1}^{n_m} (1/\delta_{l,i,j}) J_{l,j}^{syn} - (1/\gamma_{l,i}) J_l^{grow} - J_{l,i}^{sink,S} = 0 \tag{14}$$

$$J_{l,j}^{syn} + J_{l,j}^{upt,M} - J_{l,j}^{leak} - (1/\gamma_{l,j}) J_l^{grow} - J_{l,j}^{sink,M} = 0 \tag{15}$$

Here, the uptake rate of the external substrate $S_i$ by cell type $l$ follows the classical Monod equation [4]

$$J_{l,i}^{upt,S} = \frac{V_{l,i}[S_i]}{K_{l,i} + [S_i]} \qquad l = 1, 2, \cdots, n_c, \;\; i = 1, 2, \cdots, n_s \tag{16}$$

where $V_{l,i}$ is the maximum uptake rate and $K_{l,i}$ is the half-maximal substrate concentration. Similarly, the uptake rate of the external building block $M_j$ by cell type $l$ follows a modified Monod equation by including terms that account for the potential inhibitions from substrates

$$J_{l,j}^{upt,M} = \frac{V_{l,j}[M_j]}{K_{l,j} + [M_j]} \left( \prod_{i=1}^{n_s} \frac{C_{l,i,j}}{C_{l,i,j} + [S_i]} \right) \qquad l = 1, 2, \cdots, n_c, \;\; j = 1, 2, \cdots, n_m \tag{17}$$

where $V_{l,j}$ is the maximum uptake rate, $K_{l,j}$ is the half-maximal metabolite concentration in the absence of the substrates $S_i$, and $C_{l,i,j}$ is the inhibition constant. We assume that substrates are the preferred sources and can repress uptake of metabolic building blocks if both types of resources are present in the environment. One well-known example is diauxic growth of *Escherichia coli* (*E. coli*) on glucose where acetate is secreted from the beginning but consumed only after glucose exhaustion [5]. This regulatory mechanism that determines hierarchical use of resources associated with the same limiting nutrient is generally termed as "catabolite repression" and has been universally found in both prokaryotic and eukaryotic microorganisms [6].

The total influxes of substrates $J_{l,i}^{upt,S}$ and metabolites $J_{l,j}^{upt,M}$ are balanced by their consumption fluxes. First, the influxes of substrates are allocated to biosynthesis of internal metabolites. The maximum fraction reserved by each substrate $\hat{S}_{l,i}$ to produce each metabolite $\hat{M}_{l,j}$ is quantified by $\phi_{l,i,j}$ ($0 \le \sum_{j=1}^{n_m} \phi_{l,i,j} \le 1$). Since the biosynthesis reaction of a metabolite generally couples multiple substrates as reactants through fixed stoichiometry ratio, its actual flux can be modeled through the Liebig's law of minimum, which states that the rate-limiting step of a reaction is determined by the reactant with the minimum ratio of its supply level relative to its stoichiometric coefficient (i.e., demand)

$$J_{l,j}^{syn} = \min_{i=1,2,\cdots,n_s} \left( \frac{\underbrace{\phi_{l,i,j} J_{l,i}^{upt,S}}_{\text{supply}}}{\underbrace{1/\delta_{l,i,j}}_{\text{demand}}} \right) \qquad l = 1,2,\cdots,n_c, \quad j = 1,2,\cdots,n_m \tag{18}$$

Similarly, we assume, for each metabolite $\hat{M}_{l,j}$, a constant fraction ($\varphi_{l,j}$) of the influx is released back to the environment

$$J_{l,j}^{leak} = \varphi_{l,j} J_{l,j}^{syn} \tag{19}$$

Second, the remaining influxes of substrates and metabolites are supplied to biomass production. The specific growth rate of cell type $l$ can also be modeled through the Liebig's law of minimum (i.e., rate is limited by the substrate or metabolite with the minimum supply-demand ratio)

$$\begin{aligned} J_l^{grow} \quad = \quad & \underbrace{\min\left( \min_{i=1,2,\cdots,n_s} \left( \frac{\underbrace{J_{l,i}^{upt,S} - \sum_{j=1}^{n_m} (1/\delta_{l,i,j}) J_{l,j}^{syn}}_{\text{supply}}}{\underbrace{1/\gamma_{l,i}}_{\text{demand}}} \right), \min_{j=1,2,\cdots,n_m} \left( \frac{\underbrace{J_{l,j}^{syn} - J_{l,j}^{leak} + J_{l,j}^{upt,M}}_{\text{supply}}}{\underbrace{1/\gamma_{l,j}}_{\text{demand}}} \right) \right)}_{\text{basal growth rate}} \\ & \cdot \underbrace{\left( \prod_{i=1}^{n_s} \frac{I_{l,i}}{I_{l,i} + [S_i]} \right)}_{\text{inhibition due to substrate toxicity}} \cdot \underbrace{\left( \prod_{j=1}^{n_m} \frac{I_{l,j}}{I_{l,j} + [M_j]} \right)}_{\text{inhibition due to metabolite toxicity}} \qquad l = 1,2,\cdots,n_c \end{aligned} \tag{20}$$

where we take the inhibitory effects on growth for substrates and metabolites if they are also toxic and $I_{l,i}$, $I_{l,j}$ represent the half-inhibition concentrations of the substrate $\hat{S}_{l,i}$ and the metabolite $\hat{M}_{l,j}$ respectively.

Finally, the remaining substrate influxes that are neither converted to metabolites nor incorporated into biomass, and the remaining metabolite influxes that are neither leaked to the environment nor incorporated into biomass, are dumped through their sink reactions

$$J_{l,i}^{sink,S} = J_{l,i}^{upt,S} - \sum_{j=1}^{n_m} (1/\delta_{l,i,j}) J_{l,j}^{syn} - (1/\gamma_{l,i}) J_l^{grow} \tag{21}$$

$$J_{l,j}^{sink,M} = J_{l,j}^{syn} + J_{l,j}^{upt,M} - J_{l,j}^{leak} - (1/\gamma_{l,j}) J_l^{grow} \tag{22}$$

Since we take the min form of the fluxes involving substrates and metabolites (Equation (18) and (20)), only the most growth-limiting nutrient (either a substrate of a metabolite) that is in short supply is explicitly conserved.

The per-capita mortality rate of cell type $l$ is assume to be a constant

$$J_l^{death} = \eta_l \qquad l = 1, 2, \cdots, n_c \tag{23}$$

#### 1.2 Unilateral Cross-Feeding between Glucose and Acetate Specialists

##### 1.2.1 Model Specification from the General Framework

Our first application is a community of two *E. coli* mutants (CV103 and CV101) with different strategies of resource utilization [7]. Briefly, the CV103 mutant has faster glucose uptake rate than the CV101 mutant. However, it cannot utilize acetate, while CV101 can grow on acetate and co-utilize both carbon sources. By secreting acetate, CV103 creates an additional carbon-source niche for CV101 and the two mutants are thus involved in a one-way cross-feeding interaction.

A chemostat model was derived using the framework we described in Sect. 1.1

$$\frac{d[G]}{dt} = D(G_0 - [G]) - J_{1,g}^{upt} N_1 - J_{3,g}^{upt} N_3 \quad (24)$$

$$\frac{dN_1}{dt} = N_1 \left( J_1^{grow} - J_1^{death} - D \right) \quad (25)$$

$$\frac{dN_3}{dt} = N_3 \left( J_3^{grow} - J_3^{death} - D \right) \quad (26)$$

$$\frac{d[A]}{dt} = D(A_0 - [A]) + \left( J_{1,a}^{leak} - J_{1,a}^{upt} \right) N_1 + J_{3,a}^{leak} N_3 \quad (27)$$

where $D$ is the dilution rate, $G_0$ and $A_0$ are the feed medium concentrations of glucose and acetate respectively, $[G]$ and $[A]$ are their concentrations in the culture vessel respectively, $N_1$ and $N_3$ are the active population densities of CV101 and CV103 respectively.

Reaction rates ($J$'s) are described as follows. $J_{1,g}^{upt}$ and $J_{3,g}^{upt}$ are the glucose uptake rates for the two mutants

$$J_{1,g}^{upt} = \frac{V_{1,g}[G]}{K_g + [G]} \quad (28)$$

$$J_{3,g}^{upt} = \frac{V_{3,g}[G]}{K_g + [G]} \quad (29)$$

where $V_{1,g}$ and $V_{3,g}$ are the maximum glucose uptake rates for CV101 and CV103 respectively, and $K_g$ is the half-saturation glucose concentration (we assume the same value for both mutants). $J_{1,a}^{upt}$ is the acetate uptake rate for CV101

$$J_{1,a}^{upt} = \frac{V_{1,a}[A]}{K_{1,a} + [A]} \frac{C_{1,g}}{C_{1,g} + [G]} \quad (30)$$

where $V_{1,a}$ is the maximum uptake rate, $K_{1,a}$ is half-saturation acetate concentration in the absence of glucose, and $C_{1,g}$ is the inhibition constant. Since the acetyl-CoA synthetase was semi-constitutively expressed in CV101 [7], we assume that the glucose repression of acetate uptake is not fully relieved and the repression effect can be quantified by $C_{1,g}$.

The acetate production rates in both CV101 $\left(J_{1,a}^{syn}\right)$ and CV103 $\left(J_{3,a}^{syn}\right)$ cells are proportional to their corresponding glucose uptake rates

$$J_{1,a}^{syn} = \phi_a \delta_a J_{1,g}^{upt} \quad (31)$$

$$J_{3,a}^{syn} = \phi_a \delta_a J_{3,g}^{upt} \quad (32)$$

where $\phi_a$ is the fraction of glucose uptake allocated to produce acetate and $\delta_a$ is the number of acetate produced per molecule of glucose consumed (we assume both parameters share the same values between

CV101 and CV103). The acetate leakage rates by both CV101 $\left(J_{1,a}^{leak}\right)$ and CV103 $\left(J_{3,a}^{leak}\right)$ cells are then proportional to their corresponding acetate production rates

$$J_{1,a}^{leak} = \varphi_a J_{1,a}^{syn} \quad (33)$$
$$J_{3,a}^{leak} = \varphi_a J_{3,a}^{syn} \quad (34)$$

where $\varphi_a$ is the proportion of acetate that is leaked to the environment (again, we assume it has the same value between CV101 and CV103).

The per-capita growth rates for CV101 $\left(J_1^{grow}\right)$ and CV103 $\left(J_3^{grow}\right)$ are determined by the most limiting nutrient supply between acetate and the remaining glucose that is not converted to acetate

$$J_1^{grow} = \min\left(\gamma_g J_{1,g}^{upt}(1-\phi_a),\ \gamma_a\left(J_{1,a}^{syn} - J_{1,a}^{leak} + J_{1,a}^{upt}\right)\right)\frac{I_{1,a}}{I_{1,a}+[A]} \quad (35)$$
$$J_3^{grow} = \min\left(\gamma_g J_{3,g}^{upt}(1-\phi_a),\ \gamma_a\left(J_{3,a}^{syn} - J_{3,a}^{leak}\right)\right)\frac{I_{3,a}}{I_{3,a}+[A]} \quad (36)$$

Note that the flux of remaining glucose is a proxy of building blocks that cannot be synthesized from acetate. Therefore, $\gamma_a$ is the biomass yield of *E. coli* on acetate and $\gamma_g$ represents the averaged yield value for *E. coli* cells to grow on building blocks other than acetate. $I_{1,a}$ and $I_{3,a}$ are the thresholds of growth inhibition of CV101 and CV103 by acetate respectively.

Lastly, cell death is not considered in this model so that

$$J_1^{death} = J_3^{death} = 0 \quad (37)$$

##### 1.2.2 Simplifying Assumptions and Justifications

Biomass yields on glucose and acetate are similar for *E. coli* cells [8, 9]: The yield of glucose was reported to be 0.45 gDW/g glucose (equivalent to 13.5 gDW/mol Carbon) [8] and the observed per-carbon acetate yield is between 10-15 gDW/mol Carbon, depending on the acetate level [9]. It is therefore reasonable to assume that glucose and acetate are completely substitutable and all building blocks that are synthesized from glucose can also be synthesized from acetate. With this assumption, we can assume that 100% of glucose influx is directed to synthesize acetate, i.e., $\phi_a = 1$, and as a result, acetate is the only growth-limiting factor.

The schematic diagram of the simplified model in shown in Fig. 2A in the main text and its equations are described below

$$\frac{d[G]}{dt} = D(G_0 - [G]) - J_{1,g}^{upt}N_1 - J_{3,g}^{upt}N_3 \quad (38)$$
$$\frac{dN_1}{dt} = N_1\left(J_1^{grow} - D\right) \quad (39)$$
$$\frac{dN_3}{dt} = N_3\left(J_3^{grow} - D\right) \quad (40)$$
$$\frac{d[A]}{dt} = D(A_0 - [A]) + \left(\varphi_a\delta_a J_{1,g}^{upt} - J_{1,a}^{upt}\right)N_1 + \varphi_a\delta_a J_{3,g}^{upt}N_3 \quad (41)$$
$$J_{1,g}^{upt} = \frac{V_{1,g}[G]}{K_g+[G]} \quad (42)$$
$$J_{3,g}^{upt} = \frac{V_{3,g}[G]}{K_g+[G]} \quad (43)$$

$$J_{1,a}^{upt} = \frac{V_{1,a}[A]}{K_{1,a}+[A]}\frac{C_{1,g}}{C_{1,g}+[G]} \tag{44}$$

$$J_{1}^{grow} = \gamma_a\left((1-\varphi_a)\delta_a J_{1,g}^{upt} + J_{1,a}^{upt}\right)\frac{I_{1,a}}{I_{1,a}+[A]} \tag{45}$$

$$J_{3}^{grow} = \gamma_a(1-\varphi_a)\delta_a J_{3,g}^{upt}\frac{I_{3,a}}{I_{3,a}+[A]} \tag{46}$$

#### 1.3 Bilateral Cross-Feeding between Lysine and Leucine auxotrophies

##### 1.3.1 Model Specification from the General Framework

Our second application is a community of two *E. coli* single-gene knockout strains: knockouts of *lysA* and *leuA* genes resulted in two engineered strains that have auxotrophic phenotype of lysine and leucine respectively [10]. The two mutants cooperate by compensating for each other's metabolic deficiency: the lysine auxotroph (ΔK) secretes leucine that can be utilized by the leucine auxotroph (ΔL), which in return facilitate growth of the lysine auxotroph by secreting lysine to the environment.

A chemostat model was derived using the framework we developed in Sect. 1.1

$$\frac{d[G]}{dt} = D(G_0 - [G]) - J^{upt}_{\Delta k,g} N_{\Delta k} - J^{upt}_{\Delta l,g} N_{\Delta l} \tag{47}$$

$$\frac{dN_{\Delta k}}{dt} = \left(J^{grow}_{\Delta k} - J^{death}_{\Delta k} - D\right) N_{\Delta k} \tag{48}$$

$$\frac{dN_{\Delta l}}{dt} = \left(J^{grow}_{\Delta l} - J^{death}_{\Delta l} - D\right) N_{\Delta l} \tag{49}$$

$$\frac{d[K]}{dt} = D(K_0 - [K]) + J^{leak}_{\Delta l,k} N_{\Delta l} - J^{upt}_{\Delta k,k} N_{\Delta k} - J^{upt}_{\Delta l,k} N_{\Delta l} \tag{50}$$

$$\frac{d[L]}{dt} = D(L_0 - [L]) + J^{leak}_{\Delta k,l} N_{\Delta k} - J^{upt}_{\Delta k,l} N_{\Delta k} - J^{upt}_{\Delta l,l} N_{\Delta l} \tag{51}$$

where $D$ is the dilution rate, $G_0$, $K_0$ and $L_0$ are the feed medium concentrations of glucose, lysine and leucine respectively, $[G]$, $[K]$ and $[L]$ are their concentrations in the culture vessel respectively, $N_{\Delta k}$ and $N_{\Delta l}$ are the population densities of active cells of the lysine and leucine auxotroph respectively.

Reaction rates ($J$'s) are described as follows. Since the two *E. coli* auxotrophies are identical except that each carries a knockout of a single different gene, they were assume to have equal glucose uptake kinetics, i.e.,

$$J^{upt}_{\Delta k,g} = J^{upt}_{\Delta l,g} = \frac{V_g[G]}{K_g + [G]} \tag{52}$$

where $V_g$ is the maximum uptake rate and $K_g$ is the half-saturation glucose concentration. $J_{\Delta k,l}$, $J_{\Delta l,k}$, $J_{\Delta k,k}$ and $J_{\Delta l,l}$ are the leucine uptake rate by the lysine auxotroph, the lysine uptake rate by the leucine auxotroph, the lysine uptake rate by the lysine auxotroph, and the leucine uptake rate of the leucine auxotroph respectively

$$J^{upt}_{\Delta k,k} = \frac{V_{\Delta k,k}[K]}{K_{\Delta k,k} + [K]} \tag{53}$$

$$J^{upt}_{\Delta l,l} = \frac{V_{\Delta l,l}[L]}{K_{\Delta l,l} + [L]} \tag{54}$$

$$J^{upt}_{\Delta l,k} = \frac{V_{\Delta l,k}[K]}{K_{\Delta l,k} + [K]} \tag{55}$$

$$J^{upt}_{\Delta k,l} = \frac{V_{\Delta k,l}[L]}{K_{\Delta k,l} + [L]} \tag{56}$$

where $V_{\Delta k,k}$, $V_{\Delta l,l}$, $V_{\Delta l,k}$, and $V_{\Delta k,l}$ are the maximum uptake rates and $K_{\Delta k,k}$, $K_{\Delta l,l}$, $K_{\Delta l,k}$ and $K_{\Delta k,l}$ are the half-saturation constants. We assume negligible inhibitory effects of glucose on amino acids uptake since no strong evidence has been found in literature.

$J^{leak}_{\Delta l,k}$ and $J^{leak}_{\Delta k,l}$ are the lysine leakage rate of the leucine auxotroph and the leucine leakage rate of the lysine auxotroph respectively

$$J^{leak}_{\Delta l,k} = \varphi_{\Delta l,k} J^{syn}_{\Delta l,k} \quad (57)$$
$$J^{leak}_{\Delta k,l} = \varphi_{\Delta k,l} J^{syn}_{\Delta k,l} \quad (58)$$

where $\varphi_{\Delta l,k}$ and $\varphi_{\Delta k,l}$ represent the proportions of lysine and leucine released back to the environment by the leucine and lysine auxotroph respectively. The biosynthesis rate of internal lysine by the leucine auxotroph $\left(J^{syn}_{\Delta l,k}\right)$ and that of internal leucine by the lysine auxotroph $\left(J^{syn}_{\Delta k,l}\right)$ are proportional to their corresponding glucose uptake rates

$$J^{syn}_{\Delta l,k} = \phi_{\Delta l,k} \delta_k J^{upt}_{\Delta l,g} \quad (59)$$
$$J^{syn}_{\Delta k,l} = \phi_{\Delta k,l} \delta_l J^{upt}_{\Delta k,g} \quad (60)$$

where $\phi_{\Delta l,k}$ and $\phi_{\Delta k,l}$ are the fractions of glucose influx allocated to produce lysine by the leucine auxotroph and leucine by the lysine auxotroph respectively, and $\delta_k$ and $\delta_l$ are the number of lysine and leucine molecules produced per molecule of glucose consumed respectively.

The per-capita growth rate of each auxotroph is determined by the most limiting factor between the auxotrophic amino acid (i.e., leucine for the leucine auxotroph and lysine for the lysine auxotroph), the non-auxotrophic amino acid (i.e., leucine for the lysine auxotroph and lysine for the leucine auxotroph), and the remaining glucose that is not converted to the non-auxotrophic amino acid

$$J^{grow}_{\Delta k} = \min\left(\gamma_g (1 - \phi_{\Delta k,l}) J^{upt}_{\Delta k,g},\ \gamma_k J^{upt}_{\Delta k,k},\ \gamma_l (J^{upt}_{\Delta k,l} + J^{syn}_{\Delta k,l} - J^{leak}_{\Delta k,l})\right) \quad (61)$$
$$J^{grow}_{\Delta l} = \min\left(\gamma_g (1 - \phi_{\Delta l,k}) J^{upt}_{\Delta l,g},\ \gamma_l J^{upt}_{\Delta l,l},\ \gamma_k (J^{upt}_{\Delta l,k} + J^{syn}_{\Delta l,k} - J^{leak}_{\Delta l,k})\right) \quad (62)$$

Note that the flux of remaining glucose is a proxy of building blocks other than the auxotrophic and non-auxotrophic amino acids. Therefore, $\gamma_k$ and $\gamma_l$ represent the biomass yields of *E. coli* on lysine and leucine respectively, and $\gamma_g$ represents the averaged yield of other building blocks. Although excessive amounts (in mM range) of certain amino acids (including leucine) are toxic to *E. coli* [11], we assume that the growth inhibitory effects of lysine and leucine are negligible, given that their concentrations were observed to be in the range of sub-mM levels in monoculture experiments [10].

Lastly, cell mortality is modeled with a first-order kinetic rate expression

$$J^{death}_{\Delta k} = \eta_{\Delta k} \quad (63)$$
$$J^{death}_{\Delta l} = \eta_{\Delta l} \quad (64)$$

where $\eta_{\Delta k}$ and $\eta_{\Delta l}$ are the rate constants.

##### 1.3.2 Simplifying Assumptions and Justifications

The model can be simplified by two assumptions: (1) leucine or lysine does not limit growth of the auxotrophic strain that synthesizes it *de novo* (i.e., its producing strain) and (2) environment leucine or lysine is not assimilated by its producing auxotrophic strain. We justified the first assumption by considering that the lysine biosynthesis pathway in the leucine auxotroph and the leucine biosynthesis pathway in the lysine auxotroph are unperturbed such that their biosynthetic fluxes may be still tightly regulated and coordinated with fluxes of other metabolic building blocks. It is therefore reasonable to assume that lysine and leucine are as growth-limiting as other building blocks (represented by the remaining glucose flux that is not converted to the two amino acids) and never become the sole most growth-limiting factor in their producing strains.

Based on the first assumption, we can simplified the model by lumping the growth effect of lysine and leucine into the growth effect of glucose in their producing strains. As a result, growth of the lysine or leucine auxotroph only depends on the availability of the amino acid that it is auxotrophic for and the remaining glucose that is not converted to the auxotrophic amino acid. Since the consumption of lysine and leucine into biomass are implicitly modeled through glucose, their biosynthesis fluxes in the model should only include the proportion that is eventually released to the environment and thus equal to their leakage fluxes, i.e., $\varphi_{\Delta k,l} = \varphi_{\Delta l,k} = 1$.

The second assumption was justified by parameter sensitivity analysis of the model developed in Sect. 1.3.1 using Markov-Chain-Monte-Carlo algorithm. Note that the model in Sect. 1.3.1 does not take any of these assumptions. As shown in Supplementary Fig. 7, the posterior distribution of amino acid uptake rates by their producing strains are 1-2 orders of magnitude lower than the distribution of amino acid uptake rates by their non-producing strains. Specifically, the median values of their posterior distributions are $V_{\Delta k,k} = 9.65\times10^{-14}$, $V_{\Delta l,l} = 1.27\times10^{-13}$, $V_{\Delta l,k} = 5.20\times10^{-16}$, and $V_{\Delta k,l} = 1.07\times10^{-15}$. Since $V_{\Delta l,k}$ and $V_{\Delta k,l}$ are orders of magnitude smaller than $V_{\Delta k,k}$ and $V_{\Delta l,l}$, we assume $V_{\Delta l,k} = V_{\Delta k,l} = 0$ in the simplified model.

The schematic diagram of the simplified model in shown in Fig. 3A in the main text and its equations are described below

$$\frac{d[G]}{dt} = D(G_0 - [G]) - J_g^{upt}(N_{\Delta k} + N_{\Delta l}) \tag{65}$$

$$\frac{dN_{\Delta k}}{dt} = \left(J_{\Delta k}^{grow} - \eta_{\Delta k} - D\right) N_{\Delta k} \tag{66}$$

$$\frac{dN_{\Delta l}}{dt} = \left(J_{\Delta l}^{grow} - \eta_{\Delta l} - D\right) N_{\Delta l} \tag{67}$$

$$\frac{d[K]}{dt} = D(K_0 - [K]) + \phi_{\Delta l,k}\delta_k J_g^{upt} N_{\Delta l} - J_{\Delta k,k}^{upt} N_{\Delta k} \tag{68}$$

$$\frac{d[L]}{dt} = D(L_0 - [L]) + \phi_{\Delta k,l}\delta_l J_g^{upt} N_{\Delta k} - J_{\Delta l,l}^{upt} N_{\Delta l} \tag{69}$$

$$J_g^{upt} = \frac{V_g[G]}{K_g + [G]} \tag{70}$$

$$J_{\Delta k,k}^{upt} = \frac{V_{\Delta k,k}[K]}{K_{\Delta k,k} + [K]} \tag{71}$$

$$J_{\Delta l,l}^{upt} = \frac{V_{\Delta l,l}[L]}{K_{\Delta l,l} + [L]} \tag{72}$$

$$J_{\Delta k}^{grow} = \min\left(\gamma_g(1 - \phi_{\Delta k,l})J_g^{upt},\ \gamma_k J_{\Delta k,k}^{upt}\right) \tag{73}$$

$$J_{\Delta l}^{grow} = \min\left(\gamma_g(1 - \phi_{\Delta l,k})J_g^{upt},\ \gamma_l J_{\Delta l,l}^{upt}\right) \tag{74}$$

#### 1.4 Multilateral Cross-Feeding between 14 Amino Acid auxotrophies

##### 1.4.1 Model Specification from the General Framework

The third application is a community of 14 *E. coli* amino acid knockouts [12], each of which is auxotrophic for cysteine (ΔC), phenylalanine (ΔF), glycine (ΔG), histidine (ΔH), isoleucine (ΔI), lysine (ΔK), leucine (ΔL), methionine (ΔM), proline (ΔP), arginine (ΔR), serine (ΔS), threonine (ΔT), tryptophan (ΔW), and tyrosine (ΔY). For simplicity, we assume that each of the 14 strains only uptakes the amino acid that it is auxotrophic for but has the potential to secrete all other 13 amino acids to the environment. This assumption was already justified for the community of the lysine and leucine auxotroph in Sect. 1.3.2 and directly applied here. By extending our 2-auxotroph model in Sect. 1.3.2, we presented the following 14-auxotroph community model

$$\frac{d[G]}{dt} = D(G_0 - [G]) - J_g^{upt} \sum_{x \in AA} N_{\Delta x} \tag{75}$$

$$\frac{dN_{\Delta x}}{dt} = \left(J_{\Delta x}^{grow} - \eta_{\Delta x} - D\right) N_{\Delta x} \qquad x \in AA \tag{76}$$

$$\frac{d[M_z]}{dt} = D(M_{0,z} - [M_z]) + \delta_z J_g^{upt} \left( \sum_{x \in AA} \varphi_{\Delta x,z} N_{\Delta x} \right) - J_{\Delta z,z}^{upt} N_{\Delta z} \qquad z \in AA \tag{77}$$

$$AA = \{c, f, g, h, i, k, l, m, p, r, s, t, w, y\} \tag{78}$$

where $D$ is the dilution rate, $G_0$ and $M_{0,z}$ are the feed medium concentrations of glucose and amino acid $z$ respectively, $[G]$ and $[M_z]$ are their concentrations in the culture vessel respectively, $N_{\Delta x}$ ($N_{\Delta z}$) is the population density of active cells of the auxotroph $\Delta x$ ($\Delta z$), $\delta_z$ is the number of molecules of amino acid $z$ produced per glucose consumed, $\varphi_{\Delta x,z}$ is the leakage fraction of amino acid $z$ secreted by the auxotroph $\Delta x$, and $\eta_{\Delta x}$ is the mortality rate of the auxotroph $\Delta x$.

Reaction rates ($J$'s) are described as follows. We assume that all auxotrophic strains have the same kinetics of glucose uptake because they are identical except for a single different gene knockout in each strain. $J_g^{upt}$ is the glucose uptake rate for all auxotrophies

$$J_g^{upt} = \frac{V_g[G]}{K_g + [G]} \tag{79}$$

where $V_g$ is the maximum uptake rate and $K_g$ is the half-saturation glucose concentration. $J_{\Delta z,z}^{upt}$ is the uptake rate of amino acid $z$ by its auxotrophic strain

$$J_{\Delta z,z}^{upt} = \frac{V_{\Delta z,z}[M_z]}{K_{\Delta z,z} + [M_z]} \tag{80}$$

where $V_{\Delta z,z}$ is the maximum uptake rate and $K_{\Delta z,z}$ is the half-saturation concentration of amino acid $z$. The per-capita growth rate of each auxotroph is determined by the more limiting factor between the auxotrophic amino acid and the remaining glucose that is not converted to the amino acid (proxy of building blocks other than the amino acid)

$$J_{\Delta x}^{grow} = \min\left( \gamma_g J_g^{upt} \left( 1 - \sum_{z \in AA} \varphi_{\Delta x,z} \right),\ \gamma_x J_{\Delta x,x}^{upt} \right) \tag{81}$$

where $\gamma_x$ and $\gamma_g$ are the biomass yields of *E. coli* on amino acid $x$ and building blocks other than amino acid $x$ respectively.

##### 1.4.2 Simplified Pairwise Batch Co-culture Model

Here we derive the analytical solution of population density fold change in pairwise coculture between any two auxotrophic strains (e.g., $\Delta x$ and $\Delta z$). Given the following assumptions,

- All cells are active, i.e., $\eta_{\Delta x} = \eta_{\Delta z} = 0$;
- Dynamics of amino acids reach equilibrium very fast, i.e., $d[M_x]/dt = d[M_z]/dt = 0$;
- Growth of both auxotrophies are limited by the auxotrophic amino acids, i.e., $J_{\Delta x}^{grow} = \gamma_x J_{\Delta x,x}^{upt}$, $J_{\Delta z}^{grow} = \gamma_z J_{\Delta z,z}^{upt}$.

, Equations (75)-(77) can be simplified to

$$\frac{d[G]}{dt} = -J_g^{upt}(N_{\Delta x} + N_{\Delta z}) \tag{82}$$

$$\frac{dN_{\Delta x}}{dt} = \gamma_x \varphi_{\Delta z,x} \delta_x J_g^{upt} N_{\Delta z} \tag{83}$$

$$\frac{dN_{\Delta z}}{dt} = \gamma_z \varphi_{\Delta x,z} \delta_z J_g^{upt} N_{\Delta x} \tag{84}$$

, and further rewritten as

$$G(0) + \frac{N_{\Delta x}(0)}{\gamma_x \varphi_{\Delta z,x}\delta_x} + \frac{N_{\Delta z}(0)}{\gamma_z \varphi_{\Delta x,z}\delta_z} = G(t) + \frac{N_{\Delta x}(t)}{\gamma_x \varphi_{\Delta z,x}\delta_x} + \frac{N_{\Delta z}(t)}{\gamma_z \varphi_{\Delta x,z}\delta_z} \tag{85}$$

$$\frac{N_{\Delta x}(0)^2}{\gamma_x \varphi_{\Delta z,x}\delta_x} - \frac{N_{\Delta z}(0)^2}{\gamma_z \varphi_{\Delta x,z}\delta_z} = \frac{N_{\Delta x}(t)^2}{\gamma_x \varphi_{\Delta z,x}\delta_x} - \frac{N_{\Delta z}(t)^2}{\gamma_z \varphi_{\Delta x,z}\delta_z} \tag{86}$$

$G(0)$ and $G(t)$ are the glucose concentration at time 0 and time $t$ respectively. $N_{\Delta x}(0)$ ($N_{\Delta z}(0)$) and $N_{\Delta x}(t)$ ($N_{\Delta z}(t)$) are the population density of the auxotroph $\Delta x$ ($\Delta z$) at time 0 and time $t$ respectively. At any moment $t$, the population densities of the two auxotrophies are given by

$$N_{\Delta x}(t) = N_{\Delta x}(0) + \frac{-\Delta_1 + \sqrt{\Delta_1^2 + \Delta_2\Delta_3}}{\Delta_2}\gamma_x \varphi_{\Delta z,x}\delta_x \tag{87}$$

$$N_{\Delta z}(t) = N_{\Delta z}(0) + \left(G(0) - G(t) + \frac{N_{\Delta x}(0) - N_{\Delta x}(t)}{\gamma_x \varphi_{\Delta z,x}\delta_x}\right)\gamma_z \varphi_{\Delta x,z}\delta_z \tag{88}$$

where the $\Delta$'s are defined as

$$\Delta_1 = N_{\Delta x}(0) + N_{\Delta z}(0) + (G(0) - G(t))\gamma_z \varphi_{\Delta x,z}\delta_z \tag{89}$$

$$\Delta_2 = \gamma_x \varphi_{\Delta z,x}\delta_x - \gamma_z \varphi_{\Delta x,z}\delta_z \tag{90}$$

$$\Delta_3 = \left((G(0) - G(t))\gamma_z \varphi_{\Delta x,z}\delta_z + 2N_{\Delta z}(0)\right)(G(0) - G(t)) \tag{91}$$

The fold change of cell density is calculated as (final cell density)/(initial cell density), i.e., $N_{\Delta x}(\infty)/N_{\Delta x}(0)$ for the auxotroph $\Delta x$ and $N_{\Delta z}(\infty)/N_{\Delta z}(0)$ for the auxotroph $\Delta z$. Since glucose is depleted after sufficient long time, we used $G(\infty) = 0$ in these calculations.

#### 1.5 Discussions on the Proportionality Assumption for Leakage Rate

In the general modeling framework described in Sect. 1.1, we assumed that the leakage rate of a metabolite is proportional to its influx rate (proportionality assumption). To understand when the proportionality assumption is valid and how it breaks down, we leveraged our previous experiences in modeling *E. coli* resource allocation [13, 14] and developed a coarse-grained single-strain model that explicitly considers metabolite concentration by characterizing the kinetic rates of metabolite biosynthesis, its passive leakage, and its utilization for biomass under enzymatic regulations. A schematic diagram of the model is shown in Fig. 11A. We summarize the main assumptions below

- We consider three reactions in the model: biosynthesis of an internal metabolite $M$ from extracellular substrate $S$, leakage of the metabolite, and consumption of the metabolite in biomass production (reaction rate $J^{con}$). The biosynthetic reaction is governed by enzyme $E_1$ and the downstream consumption of the metabolite is mediated by enzyme $E_2$ whose activity can be inhibited by $E_2$-targeting antibiotic $A$. Metabolite leakage is assume to be passive or active with a constant enzyme level. The total protein density (i.e., amino acid concentration) of $E_1$ and $E_2$ is a constant $\alpha$;

- The $E_2$ protein production rate is proportional to the translational capacity allocated to its biosynthesis, which further equals to the total protein synthesis rate allocated to $E_1$ and $E_2$ together multiplied by the relative partitioning between the two. The total production rate of $E_1$ and $E_2$ is equal to $J^{grow}\alpha$, where $J^{grow}$ is the specific growth rate and $\alpha$, as defined above, is the concentration of total amino acids contained in $E_1$ and $E_2$. This is because the specific growth rate in balanced growth is defined as $(dX/dt)/X$, i.e., the mass production rate of any biological component $X$ per mass of $X$. When $X$ represents the total number of amino acids (as an approximation of mass) in $E_1$ and $E_2$, specific growth rate is then equal to the amino acid production rate of $E_1$ and $E_2$ per total amino acids in $E_1$ and $E_2$ (i.e., $\alpha$). Therefore, the amino acid production rate of $E_1$ and $E_2$ is equal to the specific growth rate $J^{grow}$ multiplied by $\alpha$;

- The relative partitioning of the translational resources between $E_1$ and $E_2$ proteins is mediated by a transcriptional regulator $R$, whose biosynthesis rate is assume to be inversely proportional to the internal metabolite concentration $[M]$. Higher $R$ concentration leads to decreased allocation of translational resources to $E_2$ and, concomitantly, increased allocation to $E_1$. The underlying logic of this regulatory architecture is that, shortage of the internal metabolite $M$ signals bacteria to reduce its consumption flux towards biomass while increasing the flux of its uptake and conversion. For example, biosynthesis of ppGpp (guanosine pentaphosphate) in response to amino acid shortage directly inhibits transcription of both ribosomal RNAs and ribosomal protein genes while promoting that of amino acid biosynthetic genes [13].

- The internal metabolite $M$ and proteins $E_1$, $E_2$ are stable in exponential phase and thus not actively degraded (but they are still subject to growth dilution). By constrast, the turnover rate of $R$ is generally much faster than the dilution rate (e.g., the half-life time of ppGpp is only 20-30 s [15]). We therefore assume first-order kinetics for its active degradation and ignore the dilution effect.

These assumptions can be translated into the following differential equations

$$\frac{d[M]}{dt} = \underbrace{\frac{k_{e,1}[E_1][S]}{K_{m,s}+[S]}}_{\substack{\text{metabolite}\\ \text{biosynthesis rate}}} - \underbrace{J^{con}}_{\substack{\text{metabolite}\\ \text{consumption rate}}} - \underbrace{J^{grow}[M]}_{\substack{\text{metabolite}\\ \text{dilution rate}}} - \underbrace{k_m([M]-M_e)}_{\substack{\text{metabolite}\\ \text{leakage rate}}} \tag{92}$$

$$\frac{d[E_2]}{dt} = \underbrace{\frac{\alpha J^{grow}}{m_{e,2}}}_{\substack{\text{maximum translational}\\ \text{capacity of } E_2}} \cdot \underbrace{\frac{K_{i,r}}{K_{i,r}+[R]}}_{\text{transcriptional regulation}} \underbrace{-J^{grow}[E_2]}_{\substack{E_2\\ \text{dilution rate}}} \tag{93}$$

$$\frac{d[R]}{dt} = \underbrace{\frac{k_r K_{m,m}}{K_{m,m}+[M]}}_{\substack{\text{transcriptional regulator}\\ \text{biosynthesis rate}}} - \underbrace{d_r[R]}_{\substack{\text{transcriptional regulator}\\ \text{degradation rate}}} \tag{94}$$

$$J^{con} = \underbrace{\frac{k_{e,2}[E_2][M]}{K_{m,m}+[M]}}_{\substack{\text{maximum metabolite}\\ \text{consumption rate}}} \cdot \underbrace{\frac{K_{i,a}}{K_{i,a}+[A]}}_{\text{antibiotic inhibition}} \tag{95}$$

$$J^{grow} = \underbrace{J^{con}}_{\substack{\text{metabolite}\\ \text{consumption rate}}} \cdot \underbrace{Y_m}_{\substack{\text{yield of}\\ \text{metabolite}}} \tag{96}$$

$$[E_1] = \underbrace{\frac{\alpha - m_{e,2}[E_2]}{m_{e,1}}}_{\substack{\text{conservation of amino acids}\\ \text{in } E_1 \text{ and } E_2}} \tag{97}$$

where $k_{e,1}$ is the maximum metabolite biosynthesis rate from substrate (depends on nutrient quality of substrate), $k_{e,2}$ is the maximum rate of metabolite consumption, $K_{m,s}$ and $K_{m,m}$ are the Michaelis constants, $k_m$ is the diffusion rate constant, $M_e$ is the concentration of metabolite in the environment, $\alpha$ is the total amino acids contained in protein $E_1$ and $E_2$, $K_{i,r}$ is the half-inhibition constant, $k_r$ is the maximum biosynthesis rate of the transcriptional regulator, $d_r$ is the first-order rate constant of its degradation, $m_{e,1}$ and $m_{e,2}$ are the number of amino acids in protein $E_1$ and $E_2$ respectively, and $Y_m$ is the yield of bacterial growth on the internal metabolite. The proportionality constant ($\varphi$) (i.e., flux ratio) between the metabolite leakage flux and its total influx is defined as

$$\varphi = \frac{k_m([M]-M_e)}{\frac{k_{e,1}[E_1][S]}{K_{m,s}+[S]}} \tag{98}$$

We consider a well-known example where $S$ represents glucose, $M$ represents amino acids, $E_1$ represents metabolic enzymes, $E_2$ represents ribosomes (ribosomal proteins), and $R$ represents ppGpp. Using parameter values listed in Supplementary Table 4, we simulated the steady state responses of the proportionality constant $\varphi$ under two types of perturbations: (1) changing the external substrate concentration $[S]$ (Supplementary Fig. 11B-E) and (2) changing the external antibiotic concentration $[A]$ (Supplementary Fig. 11F-I). We also varied the diffusion rate constant $k_m$ in the simulations. Our simulation results suggest the following

- Increasing external substrate concentration increases growth rate (Supplementary Fig. 11B) while increasing antibiotic concentration decreases growth rate (Supplementary Fig. 11F);
- Increasing external substrate concentration leads to a positive correlation between the influx of the internal metabolite and its concentration (Supplementary Fig. 11C) while increasing antibiotic concentration leads to a negative correlation (Supplementary Fig. 11G);
- Increasing external substrate concentration and antibiotic concentration both lead to higher concentration of the internal metabolite (Supplementary Fig. 11D,H) ;

- $\varphi$ remains unchanged by increasing external substrate concentration (Supplementary Fig. 11E) but it increases substantially at higher antibiotic concentration (Supplementary Fig. 11I).

From the findings above, we concluded that the proportionality assumption may be valid for an internal metabolite when its concentration was perturbed at the upstream of the metabolite (e.g., change external substrate concentration) since it couples the leakage with upstream biosynthesis. However, it is shown to break down when the perturbation is applied from the downstream of the metabolite (e.g., change ribosome-targeting antibiotic concentration). This is also expected because the proportionality assumption does not take feedback regulation from the downstream reactions and metabolites into accounts.

#### 2 Supplementary Tables

| Parameter | Unit | Default value | Literature | MCMC (Q1) | MCMC (Q2) | MCMC (Q3) | MCMC (95% CI) |
|---|---|---|---|---|---|---|---|
| $V_{1,g}$ | $mmol/h$ | $5.52\times10^{-13}$ | $2.99\times10^{-14}$ (Ref. 7) | $3.92\times10^{-13}$ | $4.64\times10^{-13}$ | $5.48\times10^{-13}$ | $[4.808,\ 4.825]\times10^{-13}$ |
| $V_{3,g}$ | $mmol/h$ | $6.28\times10^{-13}$ | $4.43\times10^{-14}$ (Ref. 7) | $4.42\times10^{-13}$ | $5.24\times10^{-13}$ | $6.17\times10^{-13}$ | $[5.427,\ 5.445]\times10^{-13}$ |
| $K_g$ | $mM$ | $1.00\times10^{-2}$ | $1.00\times10^{-2}$ (Ref. 7, *) | | | | |
| $I_{1,a}$ | $mM$ | $2.63\times10^{1}$ | $4.00\times10^{1}$ (Ref. 16) | $3.31\times10^{1}$ | $3.91\times10^{1}$ | $4.84\times10^{1}$ | $[4.263,\ 4.281]\times10^{1}$ |
| $I_{3,a}$ | $mM$ | $1.13\times10^{2}$ | $1.00\times10^{2}$ (Ref. 16) | $1.39\times10^{2}$ | $1.98\times10^{2}$ | $3.57\times10^{2}$ | $[4.642,\ 4.772]\times10^{2}$ |
| $C_{1,g}$ | $mM$ | $8.70\times10^{-1}$ | $1.18\times10^{1}$ (Ref. 17) | $3.17\times10^{-1}$ | 1.23 | 4.78 | $[1.261,\ 1.319]\times10^{1}$ |
| $V_{1,a}$ | $mmol/h$ | $4.26\times10^{-13}$ | $1.26\times10^{-13}$ (Ref. 7) | $2.51\times10^{-13}$ | $3.45\times10^{-13}$ | $5.23\times10^{-13}$ | $[4.266.\ 4.299]\times10^{-13}$ |
| $K_{1,a}$ | $mM$ | $2.00\times10^{-1}$ | $2.00\times10^{-1}$ (Ref. 7, *) | | | | |
| $\gamma_a$ | $1/mmol$ | $2.00\times10^{11}$ | $[0.67,\ 1.00]\times10^{11}$ (Ref. 9) | $1.90\times10^{11}$ | $2.31\times10^{11}$ | $2.92\times10^{11}$ | $[2.454,\ 2.463]\times10^{11}$ |
| $\varphi_a$ | | $3.30\times10^{-1}$ | | $2.98\times10^{-1}$ | $3.67\times10^{-1}$ | $4.46\times10^{-1}$ | $[3.749,\ 3.763]\times10^{-1}$ |
| $\delta_a$ | | 3.00 | Constant (*) | | | | |

**Supplementary Table 1:** Estimated parameter values for the simplified unilateral cross-feeding model (see Sect. 1.2.2 for model details). The column "Default value" are the parameter values obtained through manual fitting and used in all simulations. The column "Literature" lists values reported in literature. A literature value is marked with “*” if the corresponding parameter is constrained to be equal to this value during parameter fitting processes. The columns "MCMC (Q1)", "MCMC (Q2)", "MCMC (Q3)" and "MCMC (95% CI)" are the 25% percentile, 50% percentile (median), 75% percentile and 95% confidence interval of their posterior distributions sampled by Markov-Chain-Monte-Carlo algorithm. To convert units of $V_{1,g}$, $V_{3,g}$, $V_{1,a}$, and $\gamma_a$ from their original data, we assume $3\times10^{-13}$ g dry mass per cell. We chose $\delta_a = 3$, which conserves carbon in the production of acetate from glucose. The "carbon conservation" choice may not reflect the actual stoichiometry in cells: one molecule of glucose can also be fermented to produce 2 molecules of acetate and 2 molecules of carbon dioxide (in this case, $\delta_a = 2$). But most importantly, the different choices of $\delta_a$ would not alter model behavior because it can be absorbed into $V_{1,g}$ and $V_{3,g}$ as a scaling factor.

| Parameter | Unit | Default value | Literature | MCMC (Q1) | MCMC (Q2) | MCMC (Q3) | MCMC (95% CI) |
|---|---|---|---|---|---|---|---|
| $V_g$ | $mmol/h$ | $3.61\times10^{-12}$ | $3.61\times10^{-12}$ (Ref. 18, *) | | | | |
| $K_g$ | $mM$ | 1.75 | $7.00\times10^{-2}$ (Ref. 19) | $9.97\times10^{-1}$ | 3.00 | 6.94 | [5.388, 5.467] |
| $V_{\Delta k,k}$ | $mmol/h$ | $8.35\times10^{-14}$ | $4.83\times10^{-14}$ (Ref. 10) | $8.29\times10^{-14}$ | $8.82\times10^{-14}$ | $9.43\times10^{-14}$ | $[8.892,\ 8.902]\times10^{-13}$ |
| $K_{\Delta k,k}$ | $mM$ | $5.00\times10^{-3}$ | $5.00\times10^{-3}$ (Ref. 20, *) | | | | |
| $V_{\Delta l,l}$ | $mmol/h$ | $1.22\times10^{-13}$ | $6.60\times10^{-14}$ (Ref. 10) | $1.23\times10^{-13}$ | $1.31\times10^{-13}$ | $1.41\times10^{-13}$ | $[1.322,\ 1.323]\times10^{-13}$ |
| $K_{\Delta l,l}$ | $mM$ | $1.07\times10^{-3}$ | $1.07\times10^{-3}$ (Ref. 21, *) | | | | |
| $\gamma_g$ | $1/mmol$ | $3.00\times10^{11}$ | $3.00\times10^{11}$ (Ref. 22, *) | | | | |
| $\gamma_k$ | $1/mmol$ | $5.72\times10^{12}$ | $9.52\times10^{12}$ (Ref. 10) | $4.31\times10^{12}$ | $4.58\times10^{12}$ | $4.86\times10^{12}$ | $[4.594,\ 4.599]\times10^{12}$ |
| $\gamma_l$ | $1/mmol$ | $2.53\times10^{12}$ | $7.05\times10^{12}$ (Ref. 10) | $2.16\times10^{12}$ | $2.34\times10^{12}$ | $2.53\times10^{12}$ | $[2.351,\ 2.354]\times10^{12}$ |
| $\varphi_{\Delta k,l}$ | | $3.20\times10^{-3}$ | | $5.20\times10^{-3}$ | $6.60\times10^{-3}$ | $8.50\times10^{-3}$ | $[7.400,\ 7.500]\times10^{-3}$ |
| $\varphi_{\Delta l,k}$ | | $1.39\times10^{-2}$ | | $9.10\times10^{-3}$ | $1.13\times10^{-2}$ | $1.41\times10^{-2}$ | $[1.199,\ 1.204]\times10^{-2}$ |
| $\delta_k$ | | 1.00 | Constant (*) | | | | |
| $\delta_l$ | | 1.00 | Constant (*) | | | | |
| $\eta_{\Delta k}$ | $1/h$ | $1.00\times10^{-1}$ | $7.32\times10^{-2}$ (Ref. 10) | $1.56\times10^{-2}$ | $3.29\times10^{-2}$ | $5.61\times10^{-2}$ | $[3.850,\ 3.880]\times10^{-2}$ |
| $\eta_{\Delta l}$ | $1/h$ | $4.00\times10^{-4}$ | $1.00\times10^{-4}$ (Ref. 10) | $5.40\times10^{-3}$ | $1.41\times10^{-2}$ | $3.08\times10^{-2}$ | $[2.080,\ 2.110]\times10^{-2}$ |

**Supplementary Table 2:** Estimated parameter values for the simplified bilateral cross-feeding model (see Sect. 1.3.2 for model details). The column "Default value" are the parameter values obtained through manual fitting and used in all simulations. The column "Literature" lists values reported in literature. A literature value is marked with “*” if the corresponding parameter is constrained to be equal to this value during parameter fitting processes. The columns "MCMC (Q1)", "MCMC (Q2)", "MCMC (Q3)" and "MCMC (95% CI)" are the 25% percentile, 50% percentile (median), 75% percentile and 95% confidence interval of their posterior distributions sampled by Markov-Chain-Monte-Carlo algorithm. To convert unit of $V_g$, $V_{\Delta k,k}$, $V_{\Delta l,l}$, $\gamma_g$, $\gamma_k$ and $\gamma_l$ from original data, we assume $3\times10^{-13}$ g dry mass per cell. $K_{\Delta k,k}$ was calculated as the geometric mean of Km values of two active lysine transport systems. We chose $\delta_k = \delta_l = 1$ to conserve carbon in the production of lysine and leucine from glucose.

| Parameter | Unit | Default value | Literature |
|---|---|---|---|
| $\gamma_c$ | $1/\mu mol$ | $2.74\times10^{9}$ | $2.74\times10^{9}$ (Ref. 12, *) |
| $\gamma_f$ | $1/\mu mol$ | $1.63\times10^{10}$ | $1.63\times10^{10}$ (Ref. 12, *) |
| $\gamma_g$ | $1/\mu mol$ | $1.04\times10^{9}$ | $1.04\times10^{9}$ (Ref. 12, *) |
| $\gamma_h$ | $1/\mu mol$ | $1.94\times10^{10}$ | $1.94\times10^{10}$ (Ref. 12, *) |
| $\gamma_i$ | $1/\mu mol$ | $8.03\times10^{9}$ | $8.03\times10^{9}$ (Ref. 12, *) |
| $\gamma_k$ | $1/\mu mol$ | $5.47\times10^{9}$ | $5.47\times10^{9}$ (Ref. 12, *) |
| $\gamma_l$ | $1/\mu mol$ | $4.63\times10^{9}$ | $4.63\times10^{9}$ (Ref. 12, *) |
| $\gamma_m$ | $1/\mu mol$ | $1.47\times10^{10}$ | $1.47\times10^{10}$ (Ref. 12, *) |
| $\gamma_p$ | $1/\mu mol$ | $1.37\times10^{9}$ | $1.37\times10^{9}$ (Ref. 12, *) |
| $\gamma_r$ | $1/\mu mol$ | $6.02\times10^{9}$ | $6.02\times10^{9}$ (Ref. 12, *) |
| $\gamma_s$ | $1/\mu mol$ | $3.76\times10^{8}$ | $3.76\times10^{8}$ (Ref. 12, *) |
| $\gamma_t$ | $1/\mu mol$ | $2.01\times10^{9}$ | $2.01\times10^{9}$ (Ref. 12, *) |
| $\gamma_w$ | $1/\mu mol$ | $4.01\times10^{10}$ | $4.01\times10^{10}$ (Ref. 12, *) |
| $\gamma_y$ | $1/\mu mol$ | $1.63\times10^{10}$ | $1.63\times10^{10}$ (Ref. 12, *) |
| $V_{\Delta c,c}$ | $\mu mol/h$ | $2.10\times10^{-11}$ | $2.10\times10^{-11}$ (Ref. 12, *) |
| $V_{\Delta f,f}$ | $\mu mol/h$ | $7.00\times10^{-12}$ | $7.00\times10^{-12}$ (Ref. 12, *) |
| $V_{\Delta g,g}$ | $\mu mol/h$ | $1.02\times10^{-10}$ | $1.02\times10^{-10}$ (Ref. 12, *) |
| $V_{\Delta h,h}$ | $\mu mol/h$ | $4.00\times10^{-12}$ | $4.00\times10^{-12}$ (Ref. 12, *) |
| $V_{\Delta i,i}$ | $\mu mol/h$ | $1.15\times10^{-10}$ | $1.15\times10^{-10}$ (Ref. 21, *) |
| $V_{\Delta k,k}$ | $\mu mol/h$ | $5.00\times10^{-11}$ | $4.83\times10^{-14}$ (Ref. 10) |
| $V_{\Delta l,l}$ | $\mu mol/h$ | $1.90\times10^{-10}$ | $1.90\times10^{-10}$ (Ref. 21, *) |
| $V_{\Delta m,m}$ | $\mu mol/h$ | $1.76\times10^{-10}$ | $4.68\times10^{-11}$ (Ref. 21) |
| $V_{\Delta p,p}$ | $\mu mol/h$ | $9.60\times10^{-11}$ | $9.60\times10^{-11}$ (Ref. 12, *) |
| $V_{\Delta r,r}$ | $\mu mol/h$ | $3.20\times10^{-11}$ | $3,69\times10^{-11}$ (Ref. 20) |
| $V_{\Delta s,s}$ | $\mu mol/h$ | $2.80\times10^{-10}$ | $2.80\times10^{-10}$ (Ref. 12, *) |
| $V_{\Delta t,t}$ | $\mu mol/h$ | $2.79\times10^{-10}$ | $1.41\times10^{-10}$ (Ref. 23) |
| $V_{\Delta w,w}$ | $\mu mol/h$ | $3.00\times10^{-12}$ | $3.00\times10^{-12}$ (Ref. 12, *) |
| $V_{\Delta y,y}$ | $\mu mol/h$ | $4.00\times10^{-12}$ | $4.00\times10^{-12}$ (Ref. 12, *) |
| $K_{\Delta c,c}$ | $\mu mol$ | $4.96\times10^{-1}$ | $4.96\times10^{-1}$ (Ref. 24, *) |
| $K_{\Delta f,f}$ | $\mu mol$ | $7.20\times10^{-1}$ | $7.20\times10^{-1}$ (Ref. 21, *) |
| $K_{\Delta g,g}$ | $\mu mol$ | 3.80 | 3.80 (Ref. 21, *) |
| $K_{\Delta h,h}$ | $\mu mol$ | $2.60\times10^{-2}$ | 1.00 (Ref. 20) |
| $K_{\Delta i,i}$ | $\mu mol$ | $2.20\times10^{-1}$ | 1.22 (Ref. 21) |
| $K_{\Delta k,k}$ | $\mu mol$ | 5.00 | 5.00 (Ref. 20, *) |
| $K_{\Delta l,l}$ | $\mu mol$ | 1.07 | 1.07 (Ref. 21, *) |
| $K_{\Delta m,m}$ | $\mu mol$ | 2.27 | 2.27 (Ref. 21, *) |
| $K_{\Delta p,p}$ | $\mu mol$ | 2.00 | 2.00 (Ref. 25, *) |
| $K_{\Delta r,r}$ | $\mu mol$ | $5.00\times10^{-2}$ | $2.60\times10^{-2}$ (Ref. 20) |
| $K_{\Delta s,s}$ | $\mu mol$ | $7.50\times10^{-1}$ | 8.95 (Ref. 26) |
| $K_{\Delta t,t}$ | $\mu mol$ | $5.40\times10^{-1}$ | $3.90\times10^{-1}$ (Ref. 23) |
| $K_{\Delta w,w}$ | $\mu mol$ | $9.00\times10^{-1}$ | $9.00\times10^{-1}$ (Ref. 21, *) |
| $K_{\Delta y,y}$ | $\mu mol$ | $3.40\times10^{-1}$ | $3.40\times10^{-1}$ (Ref. 27, *) |
| $\eta_{\Delta c}$ | $1/h$ | $1.50\times10^{-1}$ | |
| $\eta_{\Delta f}$ | $1/h$ | $3.00\times10^{-1}$ | |

| | | | |
|---|---|---|---|
| $\eta_{\Delta g}$ | $1/h$ | $2.00\times10^{-1}$ | |
| $\eta_{\Delta h}$ | $1/h$ | $2.00\times10^{-1}$ | |
| $\eta_{\Delta i}$ | $1/h$ | $4.00\times10^{-1}$ | |
| $\eta_{\Delta k}$ | $1/h$ | 0.00 | $7.32\times10^{-2}$ (Ref. 10) |
| $\eta_{\Delta l}$ | $1/h$ | $2.00\times10^{-1}$ | $1.00\times10^{-4}$ (Ref. 10) |
| $\eta_{\Delta m}$ | $1/h$ | 0.00 | |
| $\eta_{\Delta p}$ | $1/h$ | $1.00\times10^{-1}$ | |
| $\eta_{\Delta r}$ | $1/h$ | 0.00 | |
| $\eta_{\Delta s}$ | $1/h$ | $7.50\times10^{-1}$ | |
| $\eta_{\Delta t}$ | $1/h$ | $2.00\times10^{-1}$ | |
| $\eta_{\Delta w}$ | $1/h$ | $5.00\times10^{-2}$ | |
| $\eta_{\Delta y}$ | $1/h$ | $1.00\times10^{-1}$ | |
| $\delta_c$ | | 2.00 | Constant (*) |
| $\delta_f$ | | 0.67 | Constant (*) |
| $\delta_g$ | | 3.00 | Constant (*) |
| $\delta_h$ | | 1.00 | Constant (*) |
| $\delta_i$ | | 1.00 | Constant (*) |
| $\delta_k$ | | 1.00 | Constant (*) |
| $\delta_l$ | | 1.00 | Constant (*) |
| $\delta_m$ | | 1.20 | Constant (*) |
| $\delta_p$ | | 1.20 | Constant (*) |
| $\delta_r$ | | 1.00 | Constant (*) |
| $\delta_s$ | | 2.00 | Constant (*) |
| $\delta_t$ | | 1.50 | Constant (*) |
| $\delta_w$ | | 0.55 | Constant (*) |
| $\delta_y$ | | 0.67 | Constant (*) |
| $V_g$ | $\mu mol/h$ | $3.61\times10^{-9}$ | $3.61\times10^{-9}$(Ref. 18, *) |
| $K_g$ | $\mu M$ | 1.75 | 1.75 (Ref. 28, *) |
| $\gamma_g$ | $1/\mu mol$ | $3.00\times10^{8}$ | $3.00\times10^{8}$ (Ref. 22, *) |

**Supplementary Table 3:** Estimated parameter values for the multilateral cross-feeding model (see Sect. 1.4.1 for model details). The default values of the amino acid leakage fractions ($\varphi_{\Delta x,z}$, $x$, $z \in \{c, f, g, h, i, k, l, m, p, r, s, t, w, y\}$) are displayed in the main text Fig. 5C. To convert unit of $V_g$ and $\gamma_g$ from original data, we assume $3\times10^{-13}$ g dry mass per cell. To convert unit of $V_{\Delta x,x}$ ($x \in \{i, k, l, m, r, t\}$) from original data, we assume $1\times10^{-12}$ g wet mass per cell and 0.2 pg protein per cell. The other $V_{\Delta x,x}$ ($x \in \{c, f, g, h, p, s, w, y\}$) values were directly estimated from the measured growth rates [12] of corresponding auxotrophies by multiplying the biomass yields of *E. coli* on these amino acids. $K_{\Delta k,k}$ and $K_{\Delta h,h}$ were calculated as the geometric mean of Km values of two active lysine and histidine transport systems respectively. The values of $\delta_x$ ($x \in \{c, f, g, h, i, k, l, m, p, r, s, t, w, y\}$) were chosen to conserve carbon in the production of amino acids from glucose.

| Parameter | Unit | Value | Source |
|---|---|---|---|
| $\alpha$ | $\mu M$ | $3.00\times10^{6}$ | Ref. 29 |
| $k_{e,1}$ | $1/h$ | $1.20\times10^{3}$ | |
| $k_{e,2}$ | $1/h$ | $7.56\times10^{4}$ | Ref. 29 |
| $m_{e,1}$ | | 325 | Ref. 19 |
| $m_{e,2}$ | | 11738 | Ref. 30 |
| $K_{m,s}$ | $\mu M$ | $1.00\times10^{2}$ | |
| $K_{m,m}$ | $\mu M$ | $2.00\times10^{1}$ | Ref. 29 |
| $K_{i,r}$ | $\mu M$ | $6.00\times10^{1}$ | |
| $k_r$ | $1/h$ | $3.60\times10^{3}$ | Ref. 29 |
| $d_r$ | $1/h$ | $1.26\times10^{2}$ | Ref. 29 |
| $Y_m$ | $1/\mu M$ | $3.33\times10^{-7}$ | |
| $M_e$ | $\mu M$ | 0.00 | |
| $K_{i,a}$ | $\mu M$ | 1.00 | |

**Supplementary Table 4:** Parameter values used in the simulation of the single-strain model described in Sect. 1.5. Note that these values are specific to *E. coli*. The value of $m_{e,2}$ was calculated by multiplying the literature value (7336 amino acids in ribosomal proteins) by a factor of 1.6 to account for tRNA-affiliated proteins. The yield coefficient of amino acid is the inverse of the total amino acid concentration of an *Escherichia coli* cell, i.e., $Y_m = 1/\alpha$. By choosing $M_e = 0$, we assume that the metabolite released to the environment does not accumulate and can be quickly metabolized by other cell types.

#### 3 Supplementary Figures

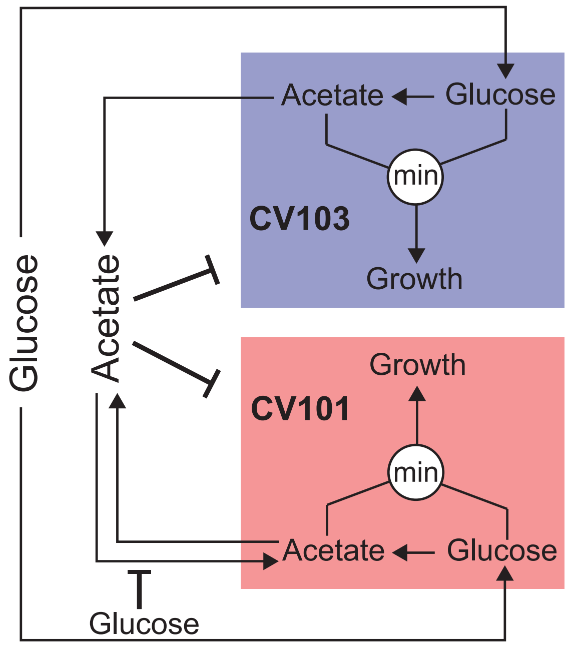

**Supplementary Figure 1:** The schematic diagram of the acetate-mediated cross-feeding model described in Sect. 1.2.1. For reference, the schematic diagram of its simplified version is shown in Fig. 2A of the main text.

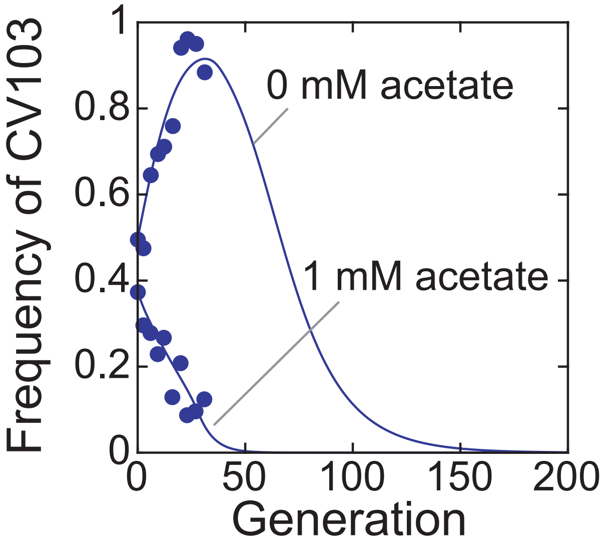

**Supplementary Figure 2:** The same as Fig. 2E of the main text but the simulated curves are shown up to 200 generations.

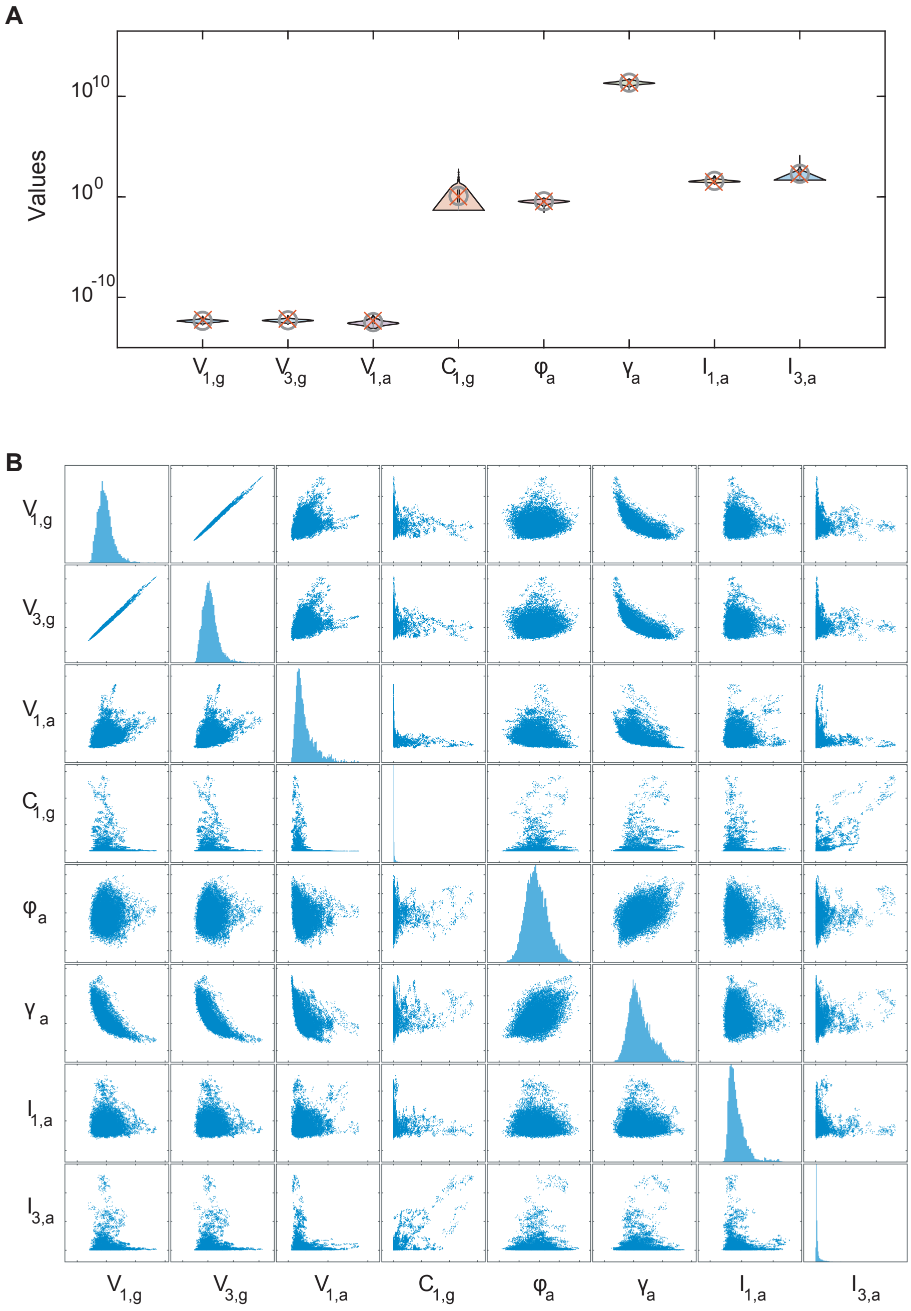

**Supplementary Figure 3:** Posterior distribution of the free parameters of the simplified acetate-mediated cross-feeding model described in Sect. 1.2.2. (A) Violin plot of the parameter distributions. Gray circles indicate the median of these distributions and red crosses indicate the values obtained through manual fitting and used in simulations. (B) Pairwise scatter plot of these distributions except that the plots along the diagonal are replaced with histograms of parameter values. Parameters not listed here are either fixed to experimentally measured values or biological constants (see Supplementary Table 1 for their values).

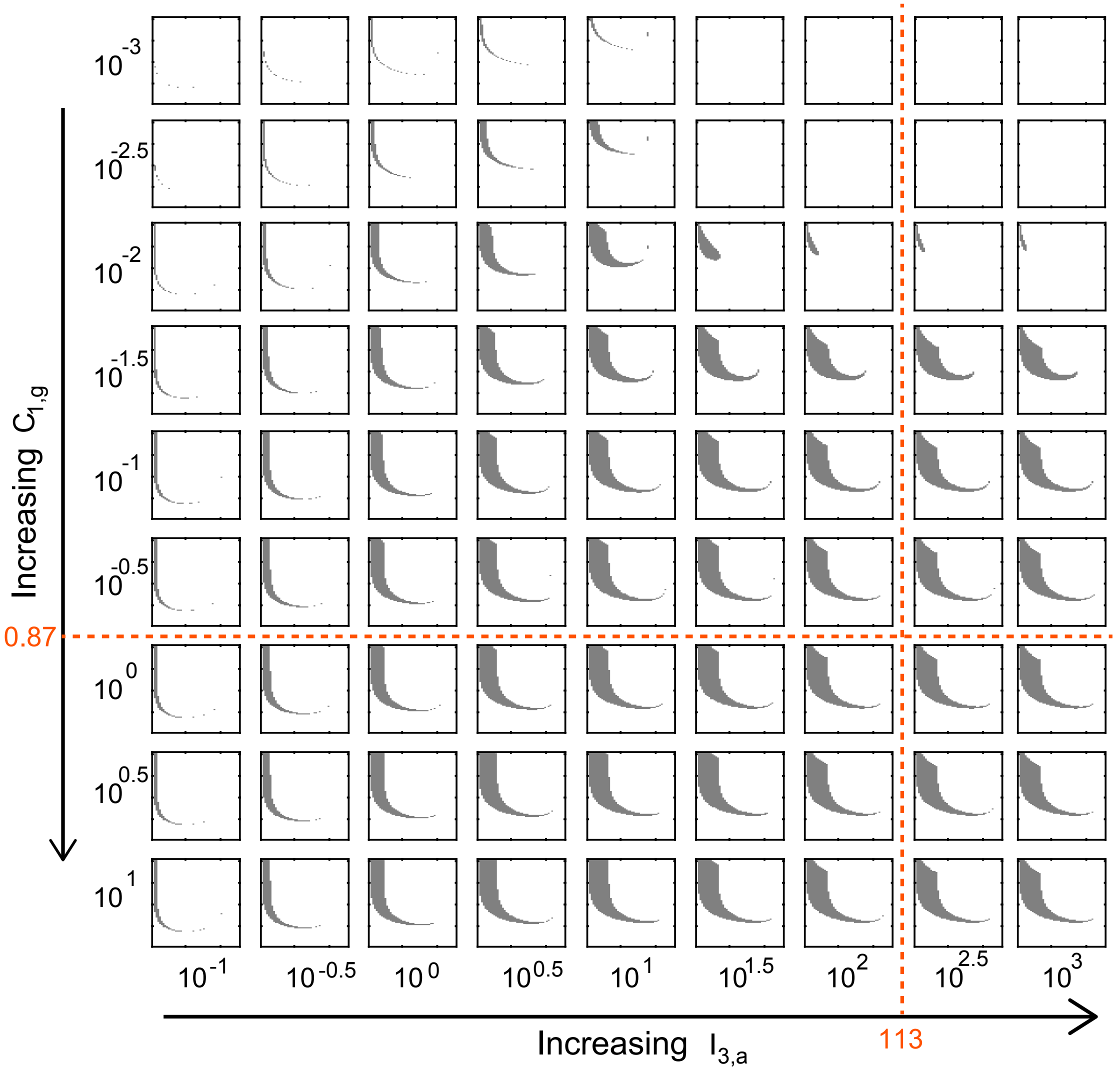

**Supplementary Figure 4:** Parameter sensitivity analysis of the coexistence region in Fig. 2G of the main text. $C_{1,g}$ and $I_{3,a}$ are the parameters that have the largest uncertainty (Supplementary Fig. 2A). Gray shading indicates the region of stable coexistence. The default values of $C_{1,g}$ and $I_{3,a}$ used to generate Fig. 2G of the main text are marked by dashed lines.

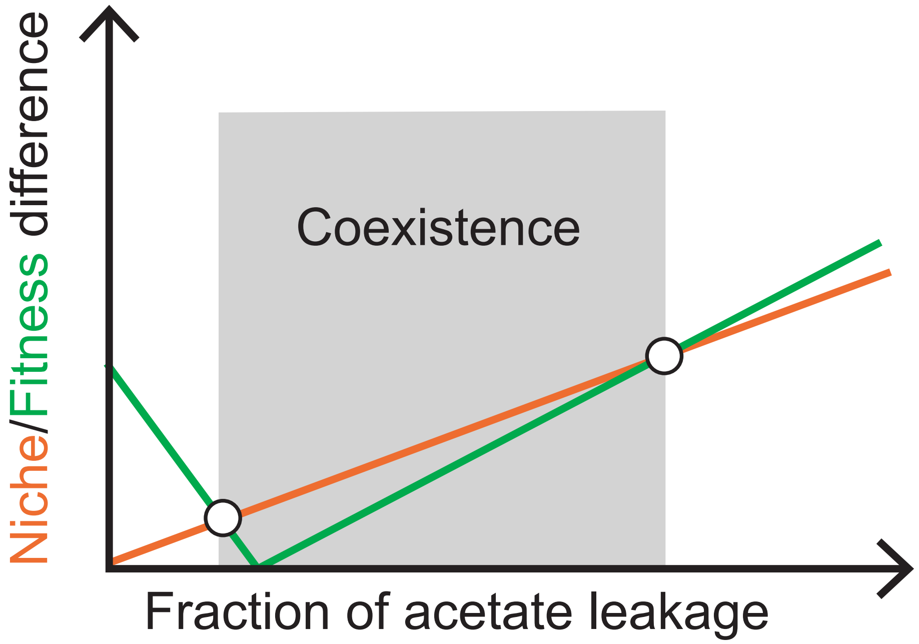

**Supplementary Figure 5:** The schematic diagram showing the qualitative changes in the niche and fitness differences with increasing proportion of acetate leakage.

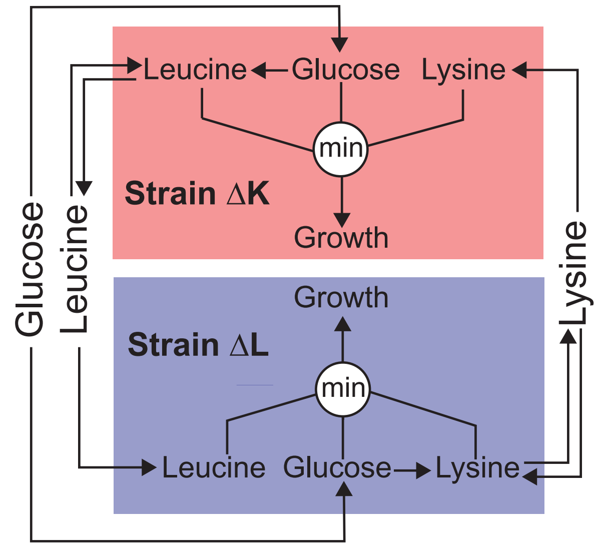

**Supplementary Figure 6:** The schematic diagram of the amino-acid-mediated cross-feeding model described in Sect. 1.3.1. For reference, the schematic diagram of its simplified version is shown in Fig. 3A of the main text.

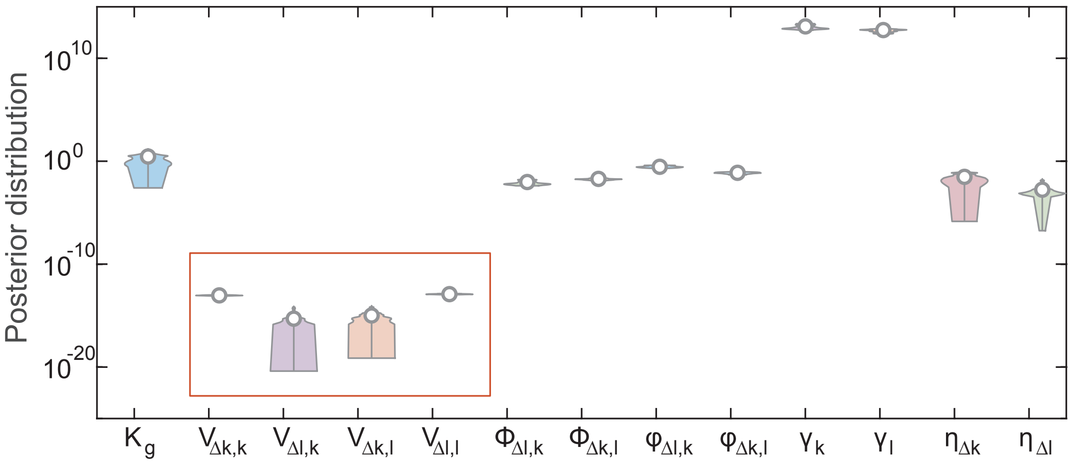

**Supplementary Figure 7:** Violin plot of the posterior distribution of the free parameters of the amino-acid-mediated cross-feeding model described in Sect. 1.3.1. Gray circles indicate the median of these distributions. The red box compares the maximum rates of amino acids uptake by their producing strains ($V_{\Delta l,k}$ and $V_{\Delta k,l}$) and those rates by their non-producing strains ($V_{\Delta k,k}$ and $V_{\Delta l,l}$). Parameters not listed here are either fixed to experimentally measured values or biological constants: $K_g = 0.01\ mM$ [7], $V_g = 3.61 \times 10^{-12}\ mmol/h$ [18], $K_{\Delta k,k} = 5.00 \times 10^{-3}\ mM$ [20], $K_{\Delta l,l} = 1.07 \times 10^{-3}\ mM$ [21], $\gamma_g = 3.00 \times 10^{11}\ 1/mmol$ [22], $\delta_k = \delta_l = 1$.

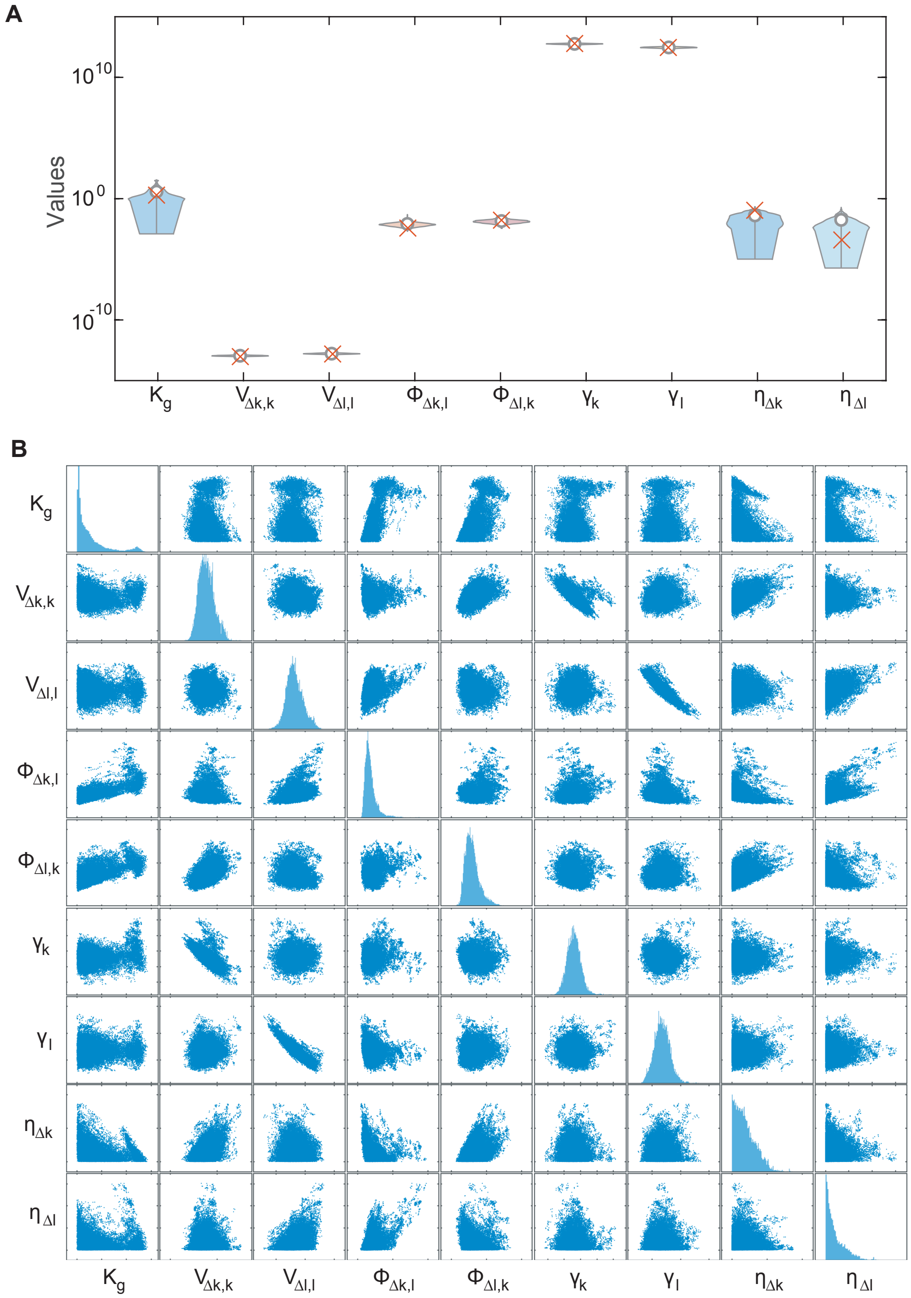

**Supplementary Figure 8:** Posterior distribution of the free parameters of the simplified amino-acid-mediated cross-feeding model described in Sect. 1.3.2. (A) Violin plot of these parameter distributions. Gray circles indicate the median of these distributions and red crosses indicate the values obtained through manual fitting and used in simulations. (B) Pairwise scatter plot of these distributions except that the plots along the diagonal are replaced with histograms of parameters values. Parameters not listed here are either fixed to experimentally measured values or biological constants (see Supplementary Table 2 for their values).

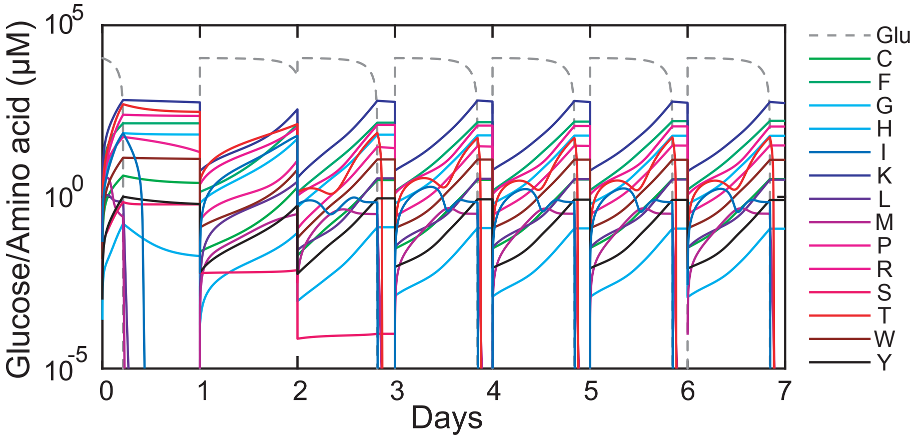

**Supplementary Figure 9:** Inferred dynamics of glucose and 14 amino acids during serial dilution of the 14-auxotroph mixture. Abbreviations: glucose (Glu), cysteine (C), phenylalanine (F), glycine (G), histidine (H), isoleucine (I), lysine (K), leucine (L), methionine (M), proline (P), arginine (R), serine (S), threonine (T), tryptophan (W), and tyrosine (Y).

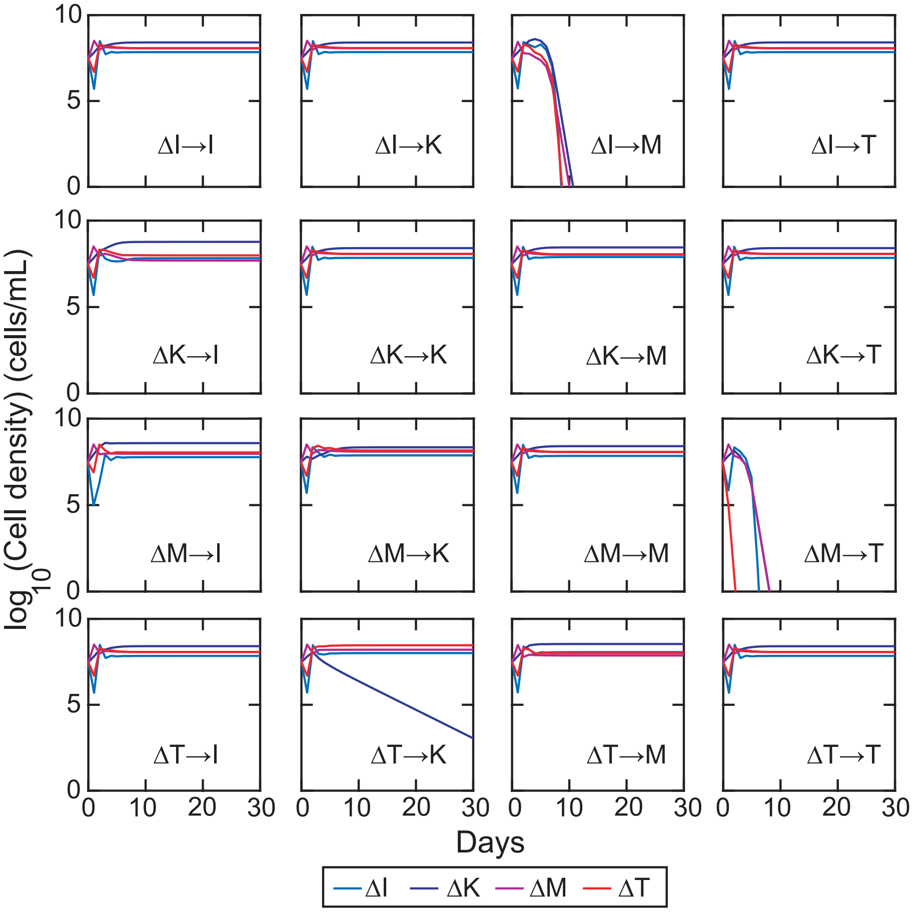

**Supplementary Figure 10:** Essentiality of amino acid secretions in stabilizing the 4-auxotroph community shown in Fig. 7A of the main text. A secretion is deemed as essential if its removal leads to strain loss. Each subplot turns off one secretion reaction: $\Delta x \to z$ indicates the secretion of amino acid $z$ by the amino acid auxotroph $\Delta x$. Our simulation results suggest that ΔI→M, ΔM→T, and ΔT→K are essential secretion fluxes. Abbreviations: isoleucine auxotroph (ΔI), lysine auxotroph (ΔK), methionine auxotroph (ΔM), threonine auxotroph (ΔT).

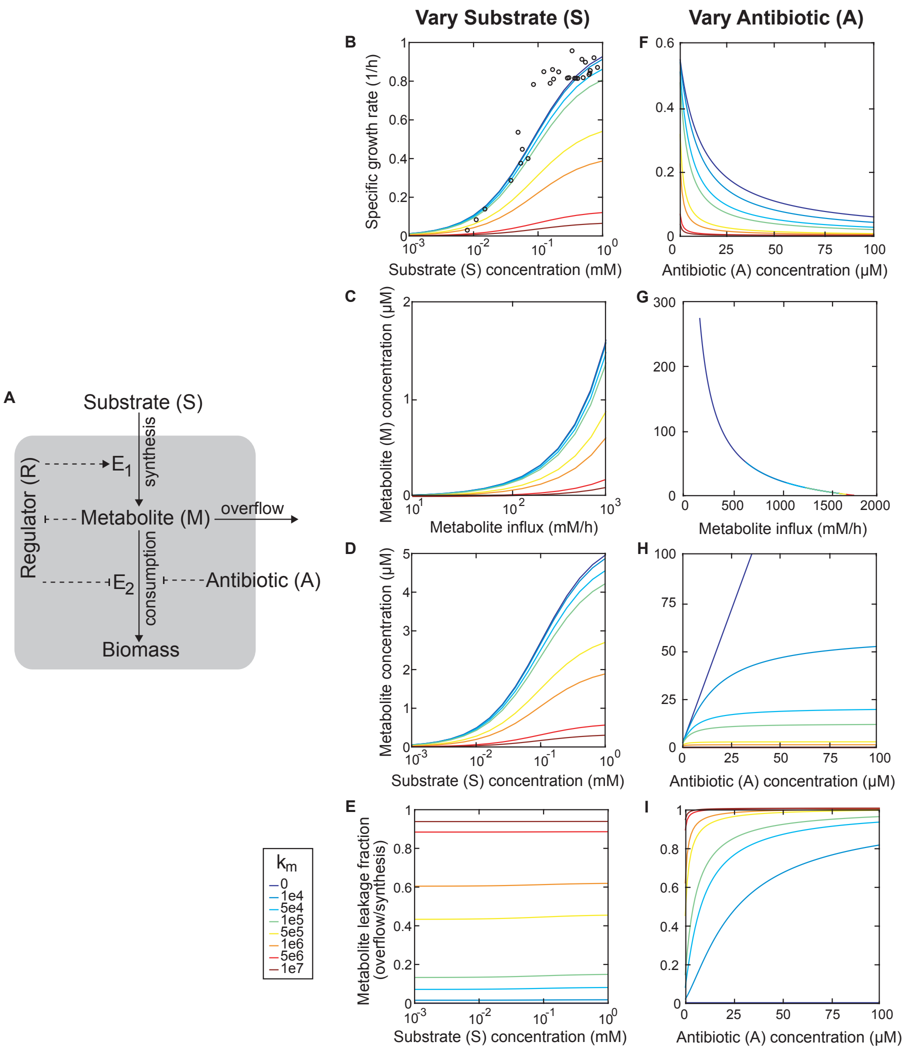

**Supplementary Figure 11:** A single-strain model of bacterial growth and metabolite overflow. (A) The schematic diagram. $E_1$ and $E_2$ are the enzymes that control biosynthesis and consumption of the metabolite $M$ respectively. Solid point arrows represent material flow. Dashed point arrows represent positive regulations and dashed blunt arrows represent negative regulations. The gray shading represents a cell. (B-E) Steady state values of various quantities by varying external substrate concentration (the antibiotic concentration is 0 $\mu M$). In particular, the model reproduces the observed Monod relationship (circles: [19]) in (B) when the diffusion rate constant ($k_m$, unit: $1/h$) is small. (F-I) Steady state values of the same quantities by varying external antibiotic concentration (the substrate concentration is 100 $\mu M$).

#### 4 Supplementary References

[1] David Tilman. *Resource competition and community structure*. Princeton university press, 1982.

[14] Chen Liao. *Integrated modeling of bacterial metabolism*. PhD thesis, University of Illinois at Urbana-Champaign, 2018.
